# TPXL-1 Activates Aurora A to Clear Contractile Ring Components from the Polar Cortex During Cytokinesis

**DOI:** 10.1101/202895

**Authors:** Sriyash Mangal, Jennifer Sacher, Taekyung Kim, Daniel Sampaio Osório, Fumio Motegi, Ana Xavier Carvalho, Karen Oegema, Esther Zanin

## Abstract

During cytokinesis, a signal from the bundled microtubules that form between the separating anaphase chromosomes promotes the accumulation of contractile ring components at the cell equator, while a signal from the centrosomal microtubule asters inhibits accumulation of contractile ring components at the cell poles. However, the molecular identity of the inhibitory signal has remained unknown. To identify molecular components of the aster-based inhibitory signal, we developed a means to monitor the removal of contractile ring proteins from the polar cortex after anaphase onset. Using this assay, we show that polar clearing is an active process that requires activation of Aurora A kinase by TPXL-1. TPXL-1 concentrates on astral microtubules coincident with polar clearing in anaphase, and its ability to recruit Aurora A and activate its kinase activity are essential for clearing. In summary, our data identify Aurora A kinase as an aster-based inhibitory signal that restricts contractile ring components to the cell equator during cytokinesis.

**SUMMARY:** During cytokinesis, centrosomal asters inhibit cortical contractility at the cell poles. Mangal et al. provide molecular insight into this phenomenon, showing that TPXL-1, which localizes to astral microtubules, activates Aurora A kinase to clear contractile ring proteins from the polar cortex.

## INTRODUCTION

Cytokinesis is the final event in mitosis that completes cell division. In animal cells, cytokinesis is accomplished by constriction of a cortical contractile ring that partitions the contents of the mother cell. The cortical accumulation of contractile ring components is controlled by the small GTPase RhoA (Piekny et al., 2005; Jordan and Canman, 2012). After anaphase onset, removal of an inhibitory Cdk1 phosphorylation recruits the RhoA guanine nucleotide exchange factor (GEF) ECT2 to the cell cortex (Su et al., 2011) and active GTP-bound RhoA and its effectors accumulate on the equatorial cortex. RhoA activates the formins, which assemble long actin filaments that make up the ring (Otomo et al., 2005), and Rho-kinase, which promotes the assembly and recruitment of myosin II (Matsumura et al., 2011). Contractile rings also include membrane-associated septin filaments and the filament cross-linker anillin, which binds to actin, septin, and myosin filaments (Piekny and Maddox, 2010; Paolo D'Avino, 2009). Recruitment of anillin to the cortex requires a direct interaction with RhoA, and, conversely, the ability to detect RhoA in an equatorial zone in fixed cells requires anillin, suggesting feedback between contractile ring assembly and RhoA localization (Piekny and Glotzer, 2008; Sun et al., 2015).

To ensure that each daughter cell inherits an equivalent genomic complement, the cortical recruitment of contractile ring components is patterned by the anaphase spindle. A signal arising in part from the spindle midzone and/or from stabilized astral microtubules oriented towards the equatorial cortex promotes the accumulation of contractile ring components on the equatorial cortex (Mishima, 2016; D’Avino et al., 2015; Green et al., 2012; von Dassow, 2009). At the same time, an inhibitory signal that antagonizes the accumulation of contractile ring proteins on the polar cortex has been proposed to arise from the centrosomal microtubule asters (Mishima, 2016; D’Avino et al., 2015; Green et al., 2012; von Dassow, 2009) and/or from kinetochores (Rodrigues et al., 2015).

Recent work in *Drosophila* sensory organ precursor and cultured human cells has suggested that kinetochore-localized protein phosphatase 1 (PP1) promotes actin clearing as the chromosomes approach the polar cortex during anaphase by dephosphorylating and inactivating ERM (ezrin/radixin/moesin) proteins (Rodrigues et al., 2015). There is also strong evidence that the centrosomal microtubule asters inhibit the accumulation of contractile ring proteins on the polar cortex. Selective disassembly of dynamic astral microtubules results in broadening of the equatorial RhoA zone/contractile ring in vertebrate cultured cells and in sea urchin embryos, suggesting that astral microtubules limit, or corral, the zone where contractile ring proteins can accumulate (Bement et al., 2005; Foe and von Dassow, 2008; von Dassow et al., 2009; Zanin et al., 2013; Murthy and Wadsworth, 2008). Consistent with this idea, laser ablation of single asters in sea urchin embryos shifts the active RhoA zone towards the ablated aster(s) (von Dassow et al., 2009). Placing microtubule asters in close proximity to the cell cortex, either genetically in *C. elegans* or mechanically in grasshopper spermatocytes, has also been shown to suppress cortical contractility in their vicinity (Chen et al., 2008; Werner et al., 2007). Despite the strong evidence for an aster-based signal that inhibits cortical contractility, its molecular identity has remained unknown.

To identify molecular components of the mechanism that prevents the accumulation of contractile ring proteins at the cell poles, we established a means to monitor removal of the contractile ring component anillin from the polar cortex during the first division of the *C. elegans* embryo. Employing this assay, we show that contractile ring proteins are actively cleared from the cell poles by a mechanism that requires TPXL-1, an Aurora A activator that localizes to astral microtubules. Activation of Aurora A by TPXL-1 is necessary for polar clearing in *C. elegans* suggesting that Aurora A mediates clearing of contractile ring components from the cell poles.

## RESULTS & DISCUSSION

### Contractile ring proteins are actively cleared from the polar cortex following anaphase onset

To explore the mechanism that antagonizes the accumulation of contractile ring proteins on the polar cortex (Fig. 1A), we monitored levels of the contractile ring protein anillin fused with GFP (GFP::anillin) at the cortex in the *C. elegans* embryo (Fig. 1C-G). We chose anillin because it mirrors the localization of active RhoA (Piekny and Glotzer, 2008), and is recruited independently of myosin II (Fig. 1D; (Maddox et al., 2005)), which allows us to monitor cortical patterning when contractility is inhibited.

We quantified cortical GFP::anillin fluorescence in central plane confocal images by performing linescans along the cortex from the anterior to the posterior pole (Fig. 1F, **Movie S1**). This analysis revealed that GFP::anillin appears on the equatorial cortex and the cortex at the anterior pole immediately following anaphase onset (see control 180s panel; Fig. 1D,G). The signal on the polar cortex then drops over the ensuing ~120s interval (between 200 and 300s after NEBD (nuclear envelope breakdown); Fig. 1D,G). In control embryos, the signal at the equator also drops after the 300s timepoint because the contractile ring constricts and moves away from the cell surface. To test whether anillin patterning depends on myosin II, we analyzed GFP::anillin after inhibiting contractility by depleting the heavy chain of myosin II (NMY-2) by RNAi. In myosin depleted embryos, the contractile ring does not constrict, and GFP::anillin levels on the equatorial cortex rose continuously over the 200s interval following anaphase onset, whereas levels on the polar cortex declined during this interval in a fashion similar to controls (Fig. 1D,G; **Movie S1**; 200-400s after NEBD). We conclude that anillin can be recruited to the equatorial cortex and cleared from the poles in the absence of myosin II based contractility.

**Figure 1:**
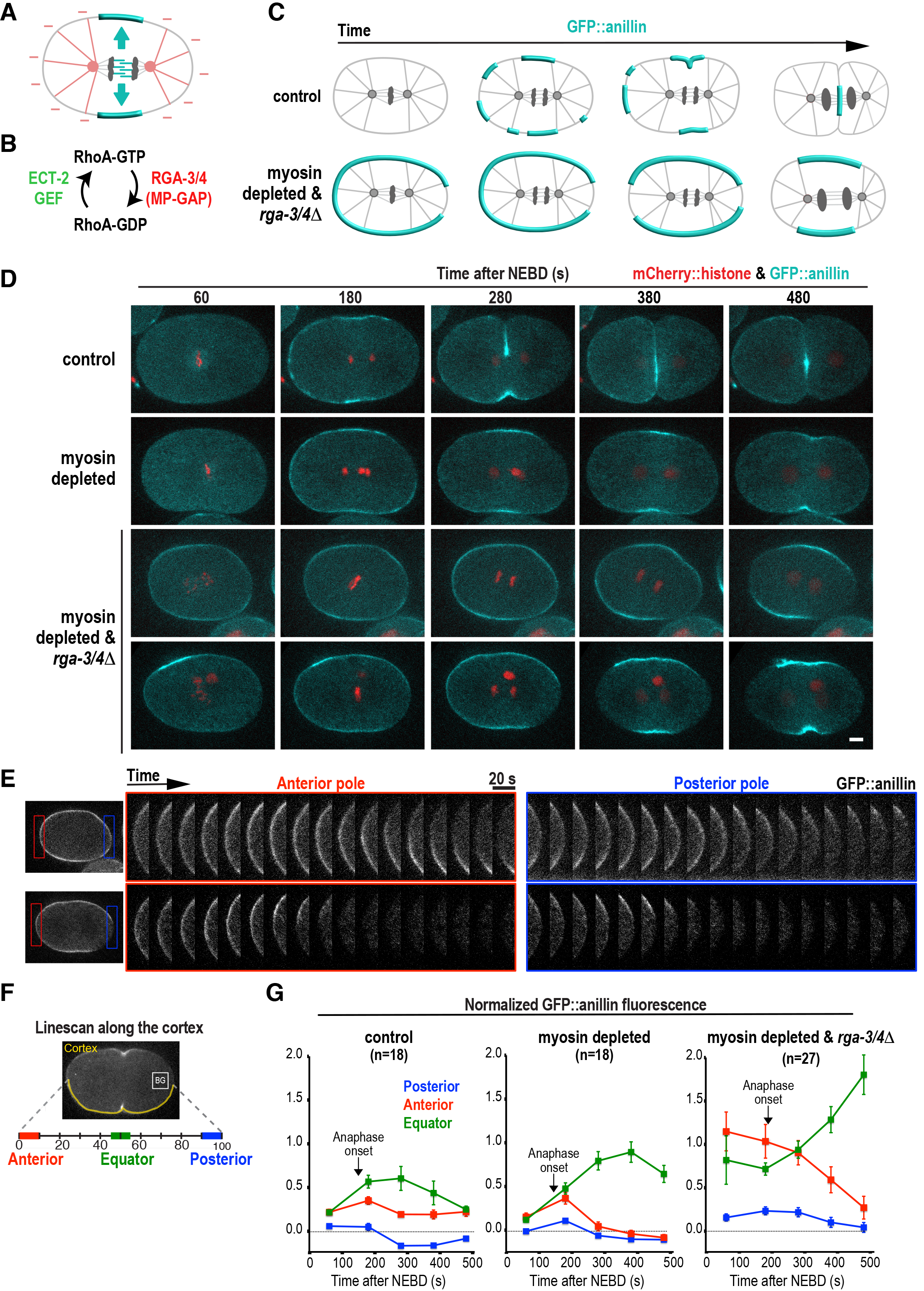
Contractile ring proteins are actively cleared from the polar cortex following anaphase onset. (**A**) Schematic illustrating the idea that the spindle provides two superimposed signals that pattern the accumulation of contractile ring proteins during cytokinesis: a stimulatory signal that promotes the accumulation of contractile ring proteins at the cell equator, and an inhibitory signal that prevents contractile ring protein accumulation at the cell poles. (**B**) During mitosis, RhoA is activated by the ECT-2 GEF and inactivated by MP-GAP (encoded by the paralogous *C. elegans* genes *rga-3* and *rga-4*). (**C**) Schematics illustrating the localization of the contractile ring marker GFP::anillin during the first mitotic division of control embryos and embryos in which contractility has been inhibited by depletion of the myosin II heavy chain NMY-2 and RhoA activity has been upregulated by deletion of *rga-3* and *rga-4* (myosin depleted & *rga-3/4*Δ). (**D**) Time-lapse confocal images of the first cell division in *C. elegans* embryos expressing GFP::anillin (***cyan***) and mCherry::histone (***red***). Single representative series are shown for control (n=9) and myosin depleted (n=9) embryos. Two representative series are shown for the myosin depleted & *rga-3/4*Δ condition; one where GFP::anillin is evenly distributed around the cortex prior to anaphase onset (***top***; n=6/14) and one where GFP::anillin is absent from the posterior cortex prior to anaphase onset (***bottom***; n=8/14). Time points are seconds after NEBD. Scale bar is 5μm. (**E**) Kymographs of the cortex at the anterior and posterior poles in the myosin depleted & *rga-3/4Δ* embryos shown in (D) beginning 180s after NEBD. (**F**) Schematic illustrating the linescan-based method used to quantify GFP::anillin fluorescence on the anterior, posterior and equatorial cortex. The mean cytoplasmic background (BG) measured in the box was subtracted, and total fluorescence was normalized by dividing by the mean intensity at the anterior pole in myosin depleted & *rga-3/4Δ* embryos 180s after NEBD. (**G**) Normalized cortical GFP::anillin fluorescence intensity is plotted for the anterior (***red***) and posterior (***blue***) poles, and for the cell equator (***green***). Plots show fluorescence versus time after NEBD in control, myosin depleted, and myosin depleted & *rga-3/4*Δ embryos. Arrows mark the average time point of anaphase onset for each condition. Error bars are SEM (standard error of the mean) and n = number of linescans.

To determine whether contractile ring proteins can be actively cleared from the polar cortex, we increased RhoA activation by eliminating the major GTPase activating protein (GAP) activity that antagonizes RhoA activation during cytokinesis (Fig. 1B). In *C. elegans*, this GAP activity is encoded by the paralogous proteins RGA-3 and RGA-4, which are homologs of MP-GAP in humans (Zanin et al., 2013; Schonegg et al., 2007; Schmutz et al., 2007). In myosin depleted & *rga-3/4*Δ embryos, GFP::anillin was already present at high levels on the equatorial and polar cortex prior to anaphase onset, allowing us to monitor how signaling by the anaphase spindle alters this pattern (Fig. 1C-G). We note that, as in control embryos, the initial accumulation of GFP::anillin on the cortex at the posterior pole was more variable than at the anterior pole (see two examples in Fig. 1D,E, **Movie S2**). We tested whether this is due to the posterior PAR proteins by depleting PAR-2 and PAR-1 in myosin depleted & *rga-3/4*Δ embryos, but this did not significantly alter GFP::anillin distribution (data not shown). Due to the variable initial accumulation of anillin on the posterior pole, we focused our analysis on comparing the dynamics of cortical GFP::anillin at the anterior pole to those at the cell equator. On the equatorial cortex in myosin depleted & *rga-3/4*Δ embryos levels of GFP::anillin were high at metaphase then increased substantially over the 300s following anaphase onset (Fig. 1D-G; 180-500s after NEBD). In contrast, at the anterior pole, levels of cortical GFP::anillin were high at metaphase and then decreased significantly over the 300s following anaphase onset (Fig. 1D-G; the same behavior was observed at the posterior pole, but with reduced magnitude due to the lower initial accumulation). These results suggest that in addition to a mechanism that promotes the accumulation of contractile ring proteins at the equator there is an active mechanism that clears contractile ring proteins from the polar cortex following anaphase onset. Due to the assay design, our results further suggest this mechanism does not require myosin-based contractility or activation of the major RhoA GAP RGA-3/4. In our subsequent analysis, we used the myosin depleted & *rga-3/4*Δ background to dissect the molecular basis for anillin clearing because the increase in the initial levels of anillin on the polar cortex due to RhoA hyperactivation facilitates analysis of its removal following anaphase onset.

### Kinetochore-localized protein phosphatase 1 (PP1) is not required for polar clearing

In *Drosophila* sensory organ precursor (SOP) cells, it has been proposed that a reduction in actin levels at the cell poles is mediated by protein phosphatase 1 (PP1) on kinetochores. In this model, cortical actin is reduced as the chromosomes and their kinetochore-associated PP1 approach the cortex (Rodrigues et al., 2015). *C. elegans* has four catalytic PP1 subunits: GSP-1, −2, −3, and −4. Since, GSP-3 and GSP-4 are only expressed during spermatogenesis (Wu et al., 2012), we focused on GSP-1 and GSP-2. Prior imaging of *in situ*-tagged GFP fusions with GSP-1 and GSP-2 revealed that they both localize transiently to kinetochores and to anaphase chromosomes (Hattersley et al., 2016; Kim et al., 2017). To test whether this chromosomal pool of PP1 is involved in polar clearing we analyzed embryos depleted of the *C. elegans* homolog of CENP-C (also called HCP-4). CENP-C is an essential component of the chromatin foundation of the kinetochore and its depletion blocks the recruitment of all outer kinetochore components (Oegema et al., 2001; Desai et al., 2003), including the two known to provide conserved PP1 docking sites, KNL-1 and MEL-28 (Espeut et al., 2012; Hattersley et al., 2016). Imaging embryos expressing GFP::GSP-1 and GFP::GSP-2 confirmed that levels of both PP1 subunits on chromosomes were greatly reduced in *hcp-4(RNAi)* embryos (Fig. 2A,B). *hcp-4(RNAi)* also blocks chromosome segregation, which would prevent any residual chromatin-associated PP1 from moving towards the poles during anaphase. Imaging myosin depleted & *rga-3/4*Δ embryos expressing a GFP-tagged centrosome marker in addition to GFP::anillin, revealed
essentially identical clearing of GFP::anillin from the polar cortex in control and *hcp-4(RNAi)* embryos (Fig. 2C-E; **Movie S3**). Cumulatively, these data suggest that the polar clearing of anillin is not mediated by kinetochore-localized PP1 in the *C. elegans* embryo. The difference from the prior analysis could be related to the fact that the kinetochores are much further away from the cortex in embryonic systems with large asters than they are during anaphase in SOP or cultured cells.

**Figure 2:**
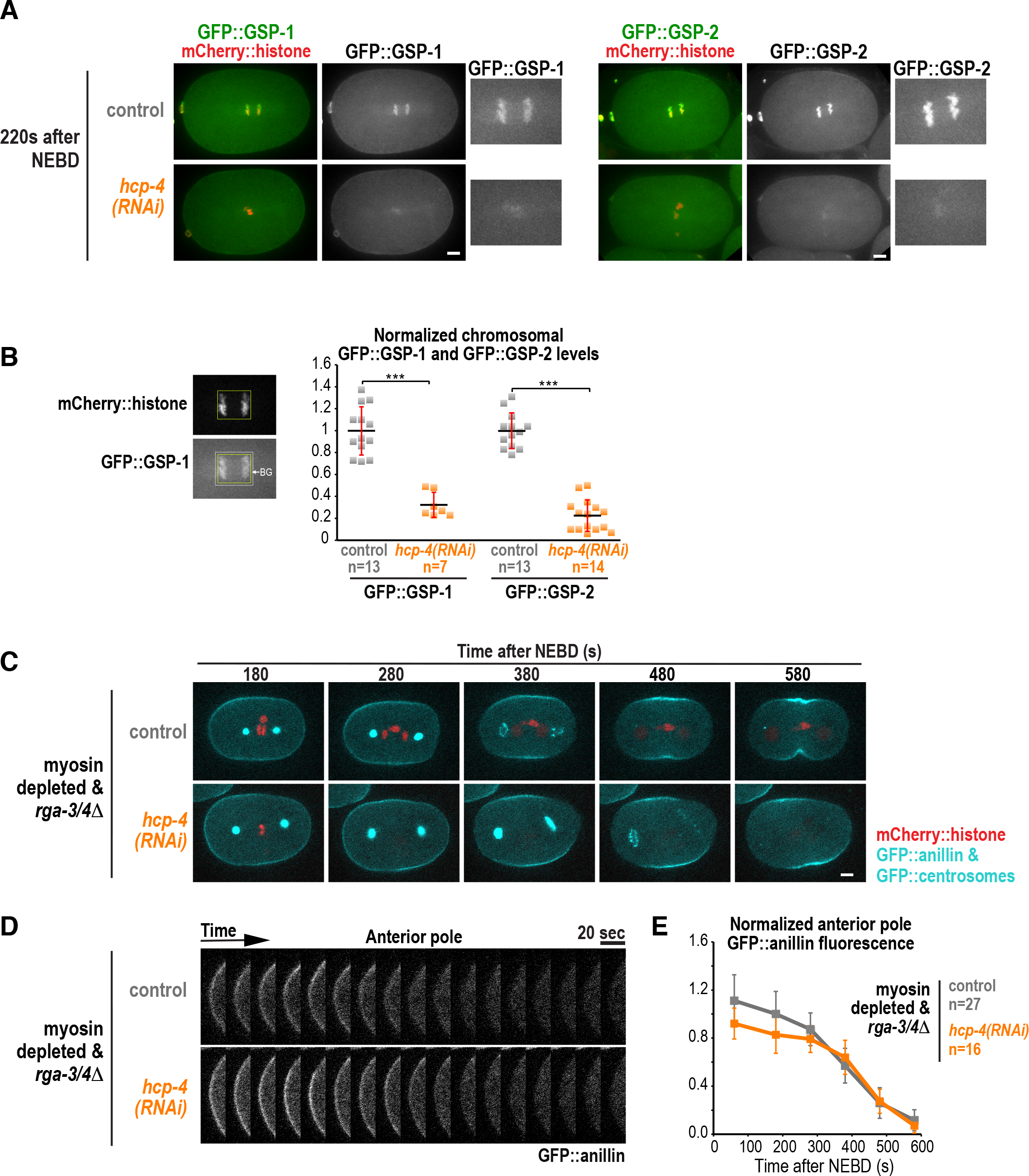
Kinetochore-localized protein phosphatase 1 (PP1) is not required for polar clearing. (**A)** Confocal images of representative control and *hcp-4(RNAi)* embryos expressing *in situ* fusions of GFP with GSP-1 (***left***) or GSP-2 (***right***) and mCherry::histone. (**B**) Mean chromosomal GFP::GSP-1 and GFP::GSP-2 fluorescence measured 200s after NEBD. n = number of embryos and error bars are SD (standard deviation). P values are obtained by student's t-test (***p<0.001). (**C**) Time-lapse confocal images of myosin depleted & *rga-3/4*Δ embryos expressing GFP::anillin, a GFP::centrosome marker (GFP::SPD-5) and mCherry::histone. A representative control embryo (***top***; n=14) and *hcp-4(RNAi)* embryo (***bottom***; n=9) are shown. (**D**) Kymographs of the cortex at the anterior pole in the embryos in (C) beginning 180s after NEBD. (**E**) Normalized cortical GFP::anillin fluorescence intensity versus time plotted for the anterior pole for the indicated conditions. n=number of linescans. Error bars are SEM and all scale bars are 5μm.

### The C. elegans TPX2 homolog, TPXL-1, is required for polar clearing

In an attempt to determine whether the proximity of the centrosomal aster is important for clearing of contractile ring proteins from the polar cortex, we analyzed embryos expressing the GFP centrosome marker and GFP::anillin that were depleted of the centrosomal Aurora A activator TPXL-1 (Ozlu et al., 2005). In TPXL-1 depleted embryos, spindles are short due to the reduced length of kinetochore microtubules, but the spindle elongates at a normal rate during anaphase (Ozlu et al., 2005). The shorter spindle length at anaphase onset in TPXL-1 depleted embryos increases the distance between the anterior centrosome and cortex during the interval when contractile ring proteins are cleared (200-400s after NEBD, Fig. 3A); this distance was increased ~2-fold 300s after NEBD in myosin depleted & *rga-3/4*Δ embryos compared to controls (21.1μm for *tpxl-1(RNAi)* versus 10.5μm in controls; Fig. 3A). Analysis of cortical GFP::anillin in the same embryos revealed that depleting TPXL-1 prevented clearing of GFP::anillin from the anterior cortex (Fig. 3B-D, **Movie S4**).

**Figure 3:**
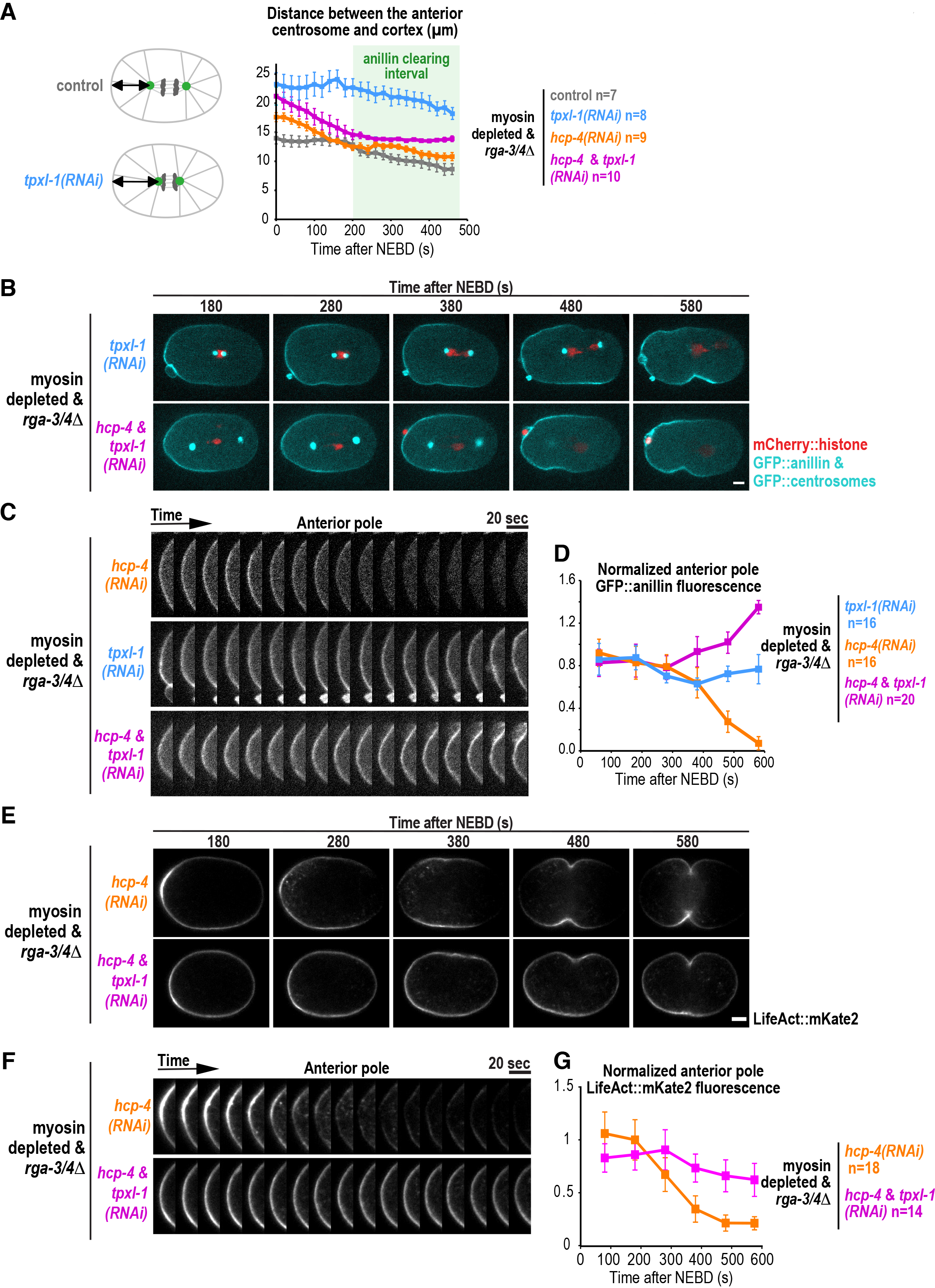
The *C. elegans* TPX2 homolog, TPXL-1, is required for polar clearing. (**A**) Schematics illustrating the effect of TPXL-1 depletion on the distance between the centrosomal aster and the cortex at the anterior pole. Graph plotting the distance between the anterior centrosome marked with GFP::SPD-5 and the anterior cell pole for the indicated conditions. n = number of embryos. (**B**) Time-lapse confocal images of myosin depleted & *rga-3/4*Δ embryos expressing GFP::anillin, a GFP::centrosome marker (***cyan***) and mCherry::histone (***red***). Representative *tpxl-1(RNAi)* (***top***; n=8) and *hcp-4 & tpxl-1(RNAi)* embryos (***bottom***; n=10) are shown. (**C**) Kymographs of the cortex at the anterior pole in the embryos shown in (B) beginning 180s after NEBD. (**D**) Normalized cortical GFP::anillin fluorescence intensity versus time plotted for the anterior pole for the indicated conditions; n = number of linescans. Images and quantification for *hcp-4(RNAi)* in C and D were reproduced from Fig. 2D and E for comparison. (**E**) Time-lapse confocal images of myosin depleted & *rga-3/4*Δ embryos expressing LifeAct::mKate2. Representative *hcp-4(RNAi)* (***top***; n=9) and *hcp-4 & tpxl-1(RNAi)* embryos (***bottom***; n=7) are shown. (**F**) Kymographs of the cortex at the anterior pole in the embryos in (E) beginning 180s after NEBD. (**G**) Normalized cortical LifeAct::mKate2 fluorescence intensity versus time plotted for the anterior pole for the indicated conditions; n = number of linescans. Error bars are SEM and scale bars are 5μm.

Depleting TPXL-1 could prevent clearing of contractile ring proteins at the cell pole because it increases the distance between the aster and the polar cortex. Alternatively, TPXL-1 could have a direct role in clearing contractile ring proteins from the polar cortex. To distinguish between these possibilities, we took advantage of the fact that disrupting kinetochore assembly by *hcp-4(RNAi)* overcomes the short kinetochore-microtubule phenotype in TPXL-1 depleted embryos and allows the asters to be separated by cortical pulling forces, restoring the centrosomal asters to a near-normal position (Ozlu et al., 2005; Lewellyn et al., 2010). Consistent with prior work, the distance between the centrosome and the anterior cortex in *hcp-4 & tpxl-1(RNAi)* embryos was comparable to the same distance in control embryos during the interval when polar clearing occurs (Fig. 3A). Despite restoration of the distance between the centrosome and polar cortex, GFP::anillin was not cleared in *hcp-4 & tpxl-1(RNAi)* embryos (Fig. 3B-D, **Movie S4**). To investigate whether filamentous actin (f-actin) is also cleared from polar cortex in TPXL-1 dependent manner we repeated the experiment in myosin depleted & *rga-3/4*Δ embryos expressing the f-actin binding probe LifeAct::mKate2. Similar to GFP::anillin, LifeAct::mKate2 was cleared from the anterior cell pole in *hcp-4(RNAi)* but not in *hcp-4 & tpxl-1 (RNAi)* embryos (Fig. 3E-G). We conclude that TPXL-1 has a direct role in promoting the clearing of anillin and f-actin from the polar cortex that is independent of its role in controlling spindle length.

To follow TPXL-1 localization we generated a functional RNAi resistant wild type TPXL-1 transgene fused to mNeonGreen (TPXL-1^WT^::NG) (Fig. S1). Consistent with prior work (Ozlu et al., 2005), imaging embryos expressing TPXL-1^WT^::NG revealed that it localizes to the periphery of the pericentriolar material in metaphase and becomes concentrated on astral microtubules in anaphase, during the interval when polar clearing is observed (Fig. 4A, **Movie S5**); a similar difference was also observed by immunofluorescence (Fig. S2A). The vertebrate TPXL-1 homolog, TPX2, has been shown to stimulate microtubule-dependent microtubule nucleation in *Xenopus* egg extracts (Petry et al., 2013). However, a previous study in metaphase *C. elegans* embryos reported that microtubule nucleation and growth rates are not affected by TPXL-1 depletion (Srayko et al., 2005; Greenan et al., 2010). To confirm these prior findings, and ensure that the failure to clear anillin from the polar cortex in TPXL-1 depleted embryos is not caused by a reduction in microtubule nucleation or in the microtubule growth rate that attenuates the microtubule asters, we analyzed the rates of microtubule nucleation and growth during the interval when clearing is observed. Microtubule nucleation and growth rates were measured using the previously described method in embryos expressing the microtubule plus end binding protein EBP-2::GFP. Consistent with prior observations (Srayko et al., 2005), we found that microtubule nucleation decreases and growth rates increase between metaphase and anaphase in control embryos (Fig. S2C-E). Microtubule nucleation was monitored as described previously; a circle with a radius of 9 μm was drawn around the anterior centrosome and kymographs were constructed along the half of the circle closest to the anterior cortex (yellow lines in Fig. S2C). Consistent with the prior work (Srayko et al., 2005), this analysis revealed that during the interval when polar clearing occurs (255-315 s after NEBD), neither the number of nucleated microtubules crossing the 9 μm radius half circle (yellow lines in Fig. 4B) nor the number of microtubules that reached the anterior cortex (blue lines in Fig. 4B) was altered by TPXL-1 depletion (Fig. 4C,D). The microtubule growth rate was also not decreased, and was instead slightly higher in *hcp-4 & tpxl-1(RNAi)* embryos compared to controls (Fig. 4E). We conclude that the failure to clear anillin from the polar cortex following anaphase in TPXL-1 depleted embryos is not caused by an attenuation of the microtubule asters due to a reduction in microtubule nucleation or in the microtubule growth rate.

**Figure 4:**
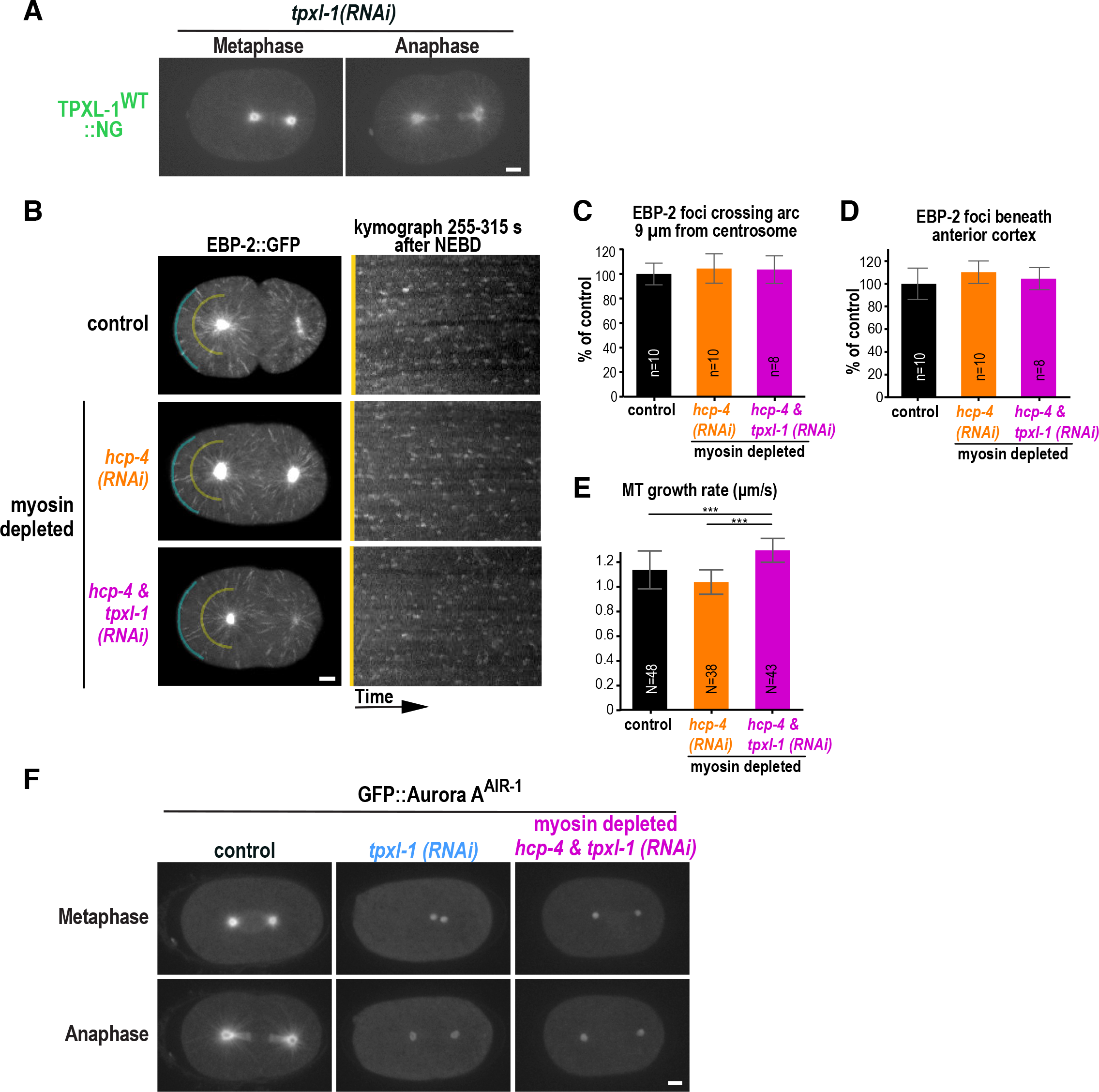
TPXL-1 recruits Aurora A^AIR-1^ to astral microtubules but is not required for astral microtubule growth or nucleation. (**A**) Confocal images of metaphase and anaphase embryos expressing TPXL-1^WT^::NG (n=8) after depletion of endogenous TPXL-1 by RNAi. (**B**) (***left***) Representative projections of the images acquired every 400 ms over a 4s interval in embryos expressing EBP-2::GFP. Images are shown for a control embryo (n=10) and for myosin depleted *hcp-4(RNAi)* (n=10) and *hcp-4 & tpxl-1(RNAi)* (n=8) embryos. EBP-2 foci were counted in kymographs made along arcs 9 μm away from the anterior centrosome (yellow, ***shown on right***) and along the anterior cortex (blue) generated for the entire 1 min movie (255-315s). (**C, D**) Graphs plot the mean number of EBP-2::GFP foci 9 μm away from the anterior centrosome (C) and beneath the anterior cortex (D) as a percentage of the mean number of foci in controls for the indicated conditions (n=number of embryos, error bars are SD). (**E**) Microtubule growth rates measured between 255-315s after NEBD for the indicated conditions (N=number of microtubules tracked in 4 or more embryos per condition and error bars are SD). (**F**) Confocal images of GFP::Aurora A^AIR^ in control (n=10), *tpxl-1(RNAi)* (n=6), and myosin depleted *hcp-4 & tpxl-1(RNAi)* (n=5) embryos. To visualize TPXL-1^WT^::NG and GFP::Aurora A^AIR^ on astral microtubules without saturating the aster centers, a gamma of 2.0 was used in (A) and 2.2 was used in (F) on images which were scaled equivalently. All scale bars are 5μm.

TPXL-1 recruits Aurora A to astral microtubules and we confirmed that Aurora A localizes to astral microtubules during anaphase in a TPXL-1 dependent fashion by imaging a GFP fusion with Aurora A (AIR-1 in *C. elegans*), (Fig. 4F, **S2B**, **Movie S6** (Ozlu et al., 2005)). Thus the failure of polar clearing in TPXL-1 depleted embryos could be due to a failure to recruit or activate Aurora A^AIR-1^ on astral microtubules.

### Polar clearing requires the ability of TPXL-1 to activate Aurora A

The vertebrate TPXL-1 homolog, TPX2 has functions that are dependent and independent of its ability to activate Aurora A kinase (Tsai et al., 2003; Eyers and Maller, 2004; Scrofani et al., 2015; Wittmann et al., 2000; Balchand et al., 2015). To determine whether the ability of TPXL-1 to promote polar clearing depends on its ability to activate Aurora A, we generated RNAi-resistant transgenes encoding wild-type TPXL-1 and a TPXL-1 mutant in which two N-terminal residues (F15 and F18) required for TPXL-1 to bind and activate Aurora A were mutated to aspartic acid (TPXL-1^FD^; (Bird and Hyman, 2008; Ozlu et al., 2005)). In addition to NG-tagged transgenes for monitoring protein localization, we also generated untagged transgenes integrated in single copy at a specific locus on chromosome II (Figs. 5A, **S1**) for functional testing. We confirmed that the transgenic proteins were expressed at levels comparable to endogenous TPXL-1 (Figs. 5B, **S1B**) and that the WT, but not the FD transgene, rescued the embryonic lethality (Fig. S1E) and the short spindle phenotype (Fig. 5D, **S1F**) resulting from endogenous TPXL-1 depletion. TPXL-1^FD^::NG localized to astral microtubules in anaphase after depletion of the endogenous protein like the WT mNeonGreen fusion (Fig. 5C), suggesting that the ability of TPXL-1 to localize to astral microtubules is independent of its ability to bind and activate Aurora A.

**Figure 5:**
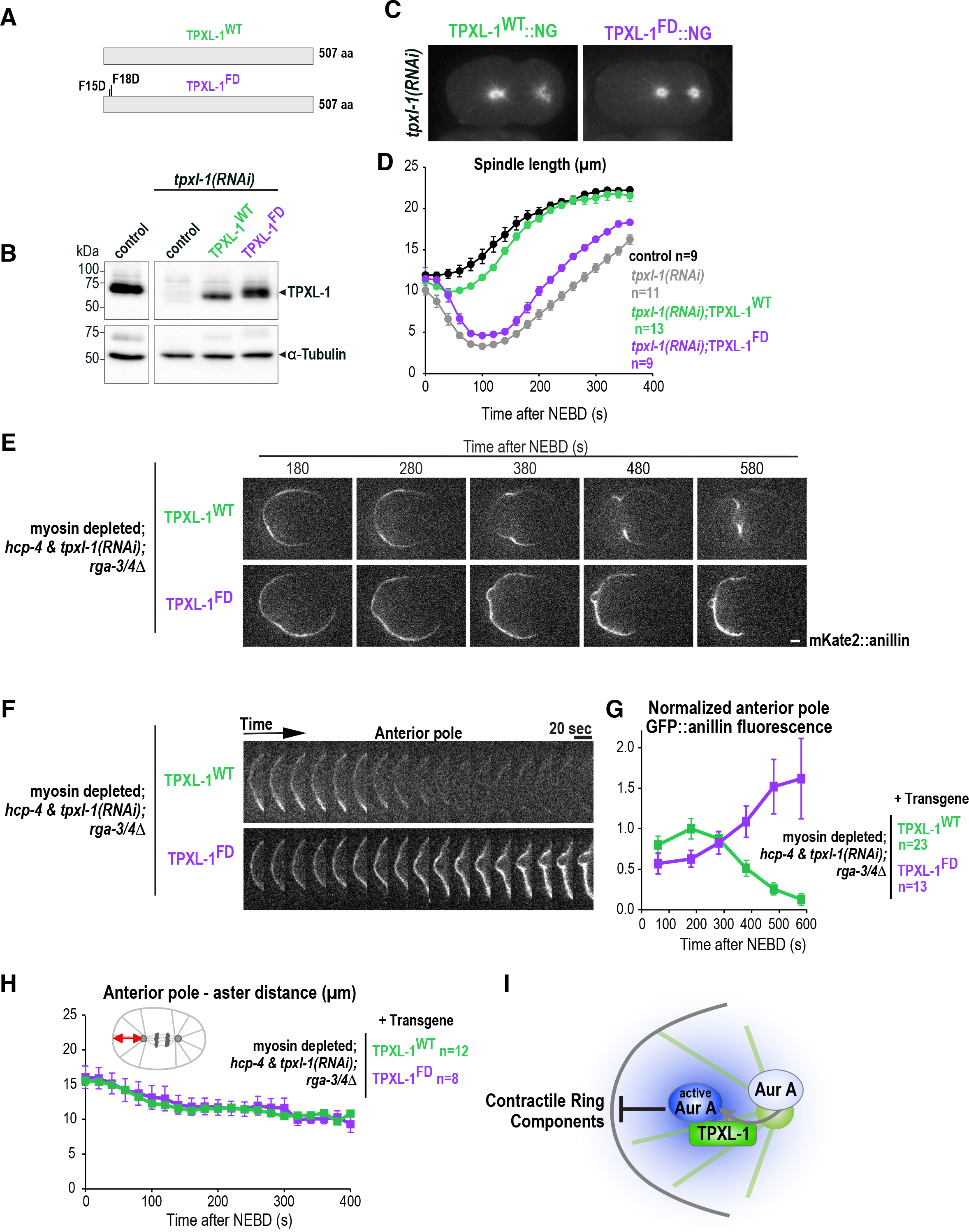
Polar clearing requires the ability of TPXL-1 to activate Aurora A. (**A**) Schematics showing the protein products of the wild-type (WT) and Aurora A binding defective (FD) *tpxl-1* transgenes. (**B**) Immunoblots of control (N2) and transgenic animals expressing RNAi-resistant TPXL-1^WT^ or TPXL-1^FD^. Endogenous TPXL-1 was depleted by RNAi as indicated and immunoblots were probed for TPXL-1 and α-tubulin as a loading control. (**C**) Confocal images of anaphase embryos expressing TPXL-1^WT^::NG (n=8) or TPXL-1^FD^::NG (n=9) after RNAi targeting the endogenous protein (**D**) Spindle length was calculated by measuring the distance between the centrosomes (Fig. S1F) over time for control (***black***), TPXL-1 depleted (*tpxl-1(RNAi)*; ***grey***) and embryos expressing TPXL-1^WT^ (***green***) or TPXL-1^FD^ (***purple***) after endogenous TPXL-1 depletion. n = number of embryos. (**E**) Time-lapse confocal images of myosin depleted & *rga-3/4*Δ embryos expressing mKate2::anillin and TPXL-1^WT^ (n=12) or TPXL-1^FD^ (n=8). Embryos were depleted of HCP-4 along with endogenous TPXL-1 to ensure comparable pole separation. (**F**) Kymographs of the cortex at the anterior pole in the embryos in (E) beginning 180s after NEBD. (**G**) Normalized cortical mKate2::anillin fluorescence intensity versus time plotted for the anterior pole for the indicated conditions; n = number of linescans. (**H**) Graph plotting the distance between the anterior aster and anterior pole for the indicated conditions. n = number of embryos. (**I**) Model illustrating how the activation of Aurora A by TPXL-1 localized to astral microtubules could generate a diffusible signal that inhibits the accumulation of contractile ring proteins on the polar cortex. All error bars are SEM and all scale bars 5μm.

To determine if the ability of TPXL-1 to activate Aurora A is required for polar clearing, we crossed the untagged WT and FD transgenes into the *rga-3/4*Δ background in the presence of a transgene expressing mKate2::anillin. Because the TPXL-1^FD^ mutant leads to a short spindle phenotype (Fig. 5D; (Ozlu et al., 2005)), we compared polar clearing in *hcp-4(RNAi)* embryos along with endogenous *tpxl-1(RNAi)* to ensure that the distance between the anterior centrosome and the polar cortex would be the same for embryos expressing the WT and FD transgenes (Fig. 5H). Whereas mKate2::anillin was cleared from the anterior cortex in the strain expressing TPXL-1^WT^, polar clearing did not occur in the strain expressing TPXL-1^FD^ (Fig. 5E-G; **Movie S7**). Thus, the ability to activate Aurora A is required for TPXL-1 to promote clearing of anillin from the polar cortex.

The current model for cytokinesis proposes that cortical contractility is patterned by two parallel signals from the mitotic spindle. A spindle midzone-dependent signal promotes contractility at the cell equator and a microtubule aster-based signal prevents contractility at the cell poles. Our work identifies TPXL-1 and Aurora A kinase as molecular components of the aster-based inhibitory signal. We demonstrate that the aster-based signal is an active signal that clears anillin and filamentous actin from the cell cortex in anaphase. Consistent with the idea that TPXL-1 and Aurora A inhibit contractility at the cell poles specifically during anaphase, we find that both become enriched on astral microtubules after anaphase onset. TPXL-1 is highly phosphorylated during mitosis (Shim et al., 2015; Wittmann et al., 2000; Heidebrecht et al., 1997) and temporal control of the association of TPXL-1 with microtubules might be achieved by the concerted action of mitotic kinases and phosphatases. Our data suggest a model in which TPXL-1 binds to astral microtubules during anaphase, where it recruits and activates Aurora A (Fig. 5I). Since Aurora A has high turnover rates (Portier et al., 2007; Stenoien et al., 2003) and is active in a gradient around monopolar spindles in Drosophila S2 cells (Ye et al., 2015) we favor a model in which active Aurora A diffuses from the astral microtubules to the adjacent cell cortex where it clears contractile ring proteins by phosphorylating specific targets. Consistent with the idea that the local activation of Aurora A can suppress cortical contractility, prior work has shown that inhibiting protein phosphatase 6 (PP6), which is the major T-loop phosphatase that negatively regulates Aurora A (Zeng et al., 2010), causes the ectopic accumulation of Aurora A on the cell cortex (Kotak et al., 2016) and globally suppresses cortical contractility (Afshar et al., 2009).

We speculate that an aster-centered Aurora A gradient together with the spindle midzone centered Aurora B activity (Fuller et al., 2008) act as landmarks of low and high contractility within the cell. Future work will be needed to determine the identity of the relevant target(s) and to determine whether Aurora A limits RhoA activation or acts on the RhoA-dependent contractile protein network.

## MATERIALS AND METHODS

### *C. elegans* strains

*C. elegans* strains used in this study are listed in Table 1. Strains were grown on NGM plates seeded with OP50 *E. coli* at 20°C according to standard procedures (Stiernagle, 2006) except OD314 and ZAN43, which were maintained at 23°C or 25°C due to silencing of the GFP::anillin encoding transgene at lower temperatures. Single copy transgenes integrated at specific chromosomal loci were generated using the MosSCI method as previously described (Frøkjær-Jensen et al., 2008). Transgenes encoding TPXL-1^WT^, TPXL-1^FD^, TPXL-1^WT^::NG and TPXL-1^FD^::NG were generated by injecting the appropriate constructs together with plasmids encoding co-injection markers (pMA122, pGH8, pCFJ190, pCFJ104) and the transposase (pCFJ601) into EG6699 to obtain single copy insertions on chromosome II. The transgenes encoding mKate2::anillin (ANI-1) or LifeAct::mKate2 were generated by injecting appropriate constructs together with the plasmids encoding transposase and co-injection markers into EG8081 to obtain a single copy insertion on chromosome IV. The transgene encoding GFP::Aurora A^AIR-1^ was integrated by microparticle bombardment.

### Cloning of expression vectors

For the expression of wild type and mNeonGreen tagged TPXL-1 (isoform A) in *C. elegans*, transgenes were cloned into pCFJ350 using Gibson cloning (NEB, E2611). Part of exon 2 and 4 and the entire exon 3 of TPXL-1 were re-encoded to render the TPXL-1 transgenes RNAi-resistant. The intron between exon 2-3 was maintained and the intron between exon 3-4 and 4-5 was removed (Fig. S1C). TPXL-1 is the second gene in an operon and therefore we used the *mex-5* promoter (488 nt) and the *tbb-2* 3’UTR (330 nt) (Zeiser et al., 2011) to drive TPXL-1 expression (Fig. S1D). Site-directed-mutagenesis was used to mutate phenylalanine 15 and 18 of TPXL-1 to aspartic acid. For expression in *C. elegans* mNeonGreen was codon-optimized using the codon adaption index (Redemann et al., 2011) and three introns were introduced. For the mKate2::anillin transgene, Gibson cloning was used to insert a sequence encoding codon-optimized mKate2 (Turek et al., 2013) into the genomic locus encoding ANI-1, including 541 nt of the *ani-1* promoter and 329 nt of its 3’ UTR and to clone this into pCFJ350. LifeAct::mKate2 transgene was generated by inserting LifeAct-mKate2 sequence in the plasmid pCFJ151 containing *mex-5* promoter and the *tbb-2* 3’UTR using Gibson cloning. To generate GFP::Aurora A transgene, Aurora A cDNA was inserted into pIC26 plasmid containing *pie-1* regulatory sequences.

### Fluorescence microscopy

For imaging *C. elegans* embryos, gravid hermaphrodites were dissected in a 4μl drop of M9 buffer on an 18×18mm coverslip and the coverslip was inverted onto a 2% agarose pad. Since *rga-3/4*Δ embryos are partially osmosensitive, all *rga-3/4*Δ mutant strains were imaged without pressure in L-15 blastomere culture medium (Zanin et al., 2013). Briefly, worms were cut open in 4μl L-15 blastomere culture medium (Edgar and Goldstein, 2012) and imaged on a 24×50mm coverslip mounted on a metal slide (Monen et al., 2005). A vaseline16 ring around the drop of L-15 blastomere culture medium was used as a spacer and a coverslip was added on top to prevent evaporation.

Images in Fig. 1, **2C, 3B, S1F** were acquired on an UltraVIEW VoX spinning disk confocal microscope (PerkinElmer) attached to an Axio Observer D1 stand (Carl Zeiss, Inc.), that was equipped with 63× 1.4 NA Plan-Apochromat oil immersion objective and 488nm and 561nm lasers. Images in Fig. 3E, **4, 5, S2C** were acquired on a Nikon eclipse Ti spinning disk confocal which was controlled by NIS Elements 4.51 and equipped with a 100× 1.45 NA Plan-Apochromat oil immersion objective, a 488nm laser line and a Hamamatsu EMCCD C9100-50 camera (1000×1000 pixel). Images in Fig. 2A were acquired at 20°C on an Andor Revolution XD Confocal System (Andor Technology) and a spinning disk confocal scanner unit (CSU-10; Yokogawa) mounted on an inverted microscope (TE2000-E; Nikon) equipped with 100x or 60× 1.4 NA Plan Apochromat lenses, and outfitted with an electron multiplication back-thinned charged-coupled device camera (iXon, Andor Technology). All immunofluorescence images were taken on a laser scanning confocal Leica TCS SP5 microscope equipped with a 63× 1.4 NA Plan-Apochromat oil immersion objective and 405nm, 488nm, 594nm lasers.

### Image quantifications

All quantification was done on raw images in Fiji (Schindelin et al., 2012). For the image analysis NEBD was defined as the time point when nucleoplasmic mCherry::histone equilibrated with the cytoplasm. In strains without a mCherry::histone marker, NEBD was defined as the time point when the border of nucleus disappeared in transmitted light images. Linescans were drawn in Fiji along the cortex from the anterior to the posterior pole at defined time points after NEBD (Fig. 1F). Sometimes the meiosis remnant resulted in a bright anterior signal at the cell cortex and these cortices were excluded. The cytoplasmic background intensity was measured in a small box close to the posterior pole and subtracted from each cortex value. The mean anterior (0-10%), equatorial (45-55%) and posterior (90-100%) fluorescence intensity was calculated for each condition and plotted over time. All anillin fluorescence intensity values were normalized to the mean anterior fluorescence intensity 180s after NEBD in myosin depleted *rga-3/4*Δ embryos. The distance between the anterior pole and anterior aster was measured by drawing a line from the center of the centrosome (GFP::SDP-5) to the anterior pole (Fig. 3A, **5H**). In 1/7 ZAN248 and 5/12 ZAN249 embryos GFP::SDP-5 was silenced and in those the transmitted light images were used. To measure the chromosomal fluorescence intensity of GFP::GSP-1 and GFP::GSP-2 (Fig. 2B) a rectangular box was drawn around the chromosome area on the mCherry::histone channel and the GFP fluorescence intensity was measured in the same box in GFP channel. Then the box was expanded by 5 pixels on each side and the fluorescence intensity was measured. The signal intensity and the area difference between the original box and the expanded box were used to calculate the background intensity per pixel, which was subtracted from the intensity of original box. Fluorescent images were processed in Fiji and Adobe Photoshop Element, graphs were plotted in Prism and Excel and figures were assembled in Affinity Designer.

### Quantification of microtubule dynamics

Microtubule nucleation rate was measured as previously published (Srayko et al., 2005). Briefly, EBP2::GFP images were acquired at 400 msec intervals for a period of 1 minute at 25°C. In Fiji, an arc (29-30 μm long) was drawn 9 μm away from the anterior centrosome and underneath the anterior polar cortex. From these arcs, kymographs were generated for 1 min time interval and EBP2::GFP dots were manually counted. To measure microtubule growth rate, individual EBP2::GFP dots emerging from the anterior centrosome were manually tracked using Fiji Manual Tracking plugin. Position of EBP2::GFP dot with respect to the centrosome was plotted at each time interval and the growth rate of individual microtubules was calculated using the slope of each line (Excel). Microtubule growth rate was analyzed in at least three embryos and 8-12 MTs were tracked for each embryo.

### Immunofluorescence

For immunostainings of *C. elegans* embryos gravid adults were cut open in 10μl of M9 on 0.1% poly-L-lysine coated glass slides. Embryos were freeze-cracked in liquid N2 and fixed for 20min in −20°C-cold 100% methanol. Slides were washed in 1xPBS (0.2% Tween) for 5min and rabbit anti-TPXL-1 (1:1000, (Ozlu et al., 2005)) or rabbit anti-Aurora A (1:200, (Hannak et al., 2001)) were incubated with mouse-anti α-tubulin antibodies (1:250, DM1-α, Sigma Aldrich) at 4°C over night. Slides were washed 3x in 1×PBS (0.2% Tween) and incubated for 1hr at room temperature with the anti-mouse Alexa488 (1:500, Molecular Probes), anti-rabbit Alexa594 (1:500, Molecular Probes), and 1μg/ml Hoechst 33258 (Sigma).

### Immunoblotting

For immunoblotting, adult worms were collected and treated as described previously (Zanin et al., 2011). Briefly, 30-40 worms were picked and washed multiple times in M9 buffer. Sample buffer was added and worms were incubated at 95°C for 5min and sonicated for 20min. Membranes were incubated with rabbit anti-TPXL-1 (1:1000, (Ozlu et al., 2005)) and mouse-anti α-tubulin (1:1000, DM1-α, Sigma Aldrich) primary antibodies and HRP-conjugated rabbit and mouse secondary antibodies (Biorad).

### *C. elegans* RNAi and lethality assays

For dsRNA production the targeting region was amplified from *C. elegans* cDNA by PCR with the T7 containing oligonucleotides listed in Table 2. The purified PCR products were used for T7 in vitro transcription (MEGAscript, Ambion). L4 or young adults were injected with dsRNA and the embryos of the injected worms were filmed 24-28hrs at 20°C or 20-24hrs at 23°C. For co-depletion experiments dsRNAs were mixed at 1:1 ratio. For *tpxl-1(RNAi)* similar depletion levels were observed after 24hrs (Fig. 5B) and 40hrs (Fig. S1B, both at 20°C). Embryonic lethality assays were performed at 20°C and injected worms were singled 24hrs and sacrificed 48hrs post injection. The number of dead embryos and larvae was counted 24hrs later.

**Table 1.**
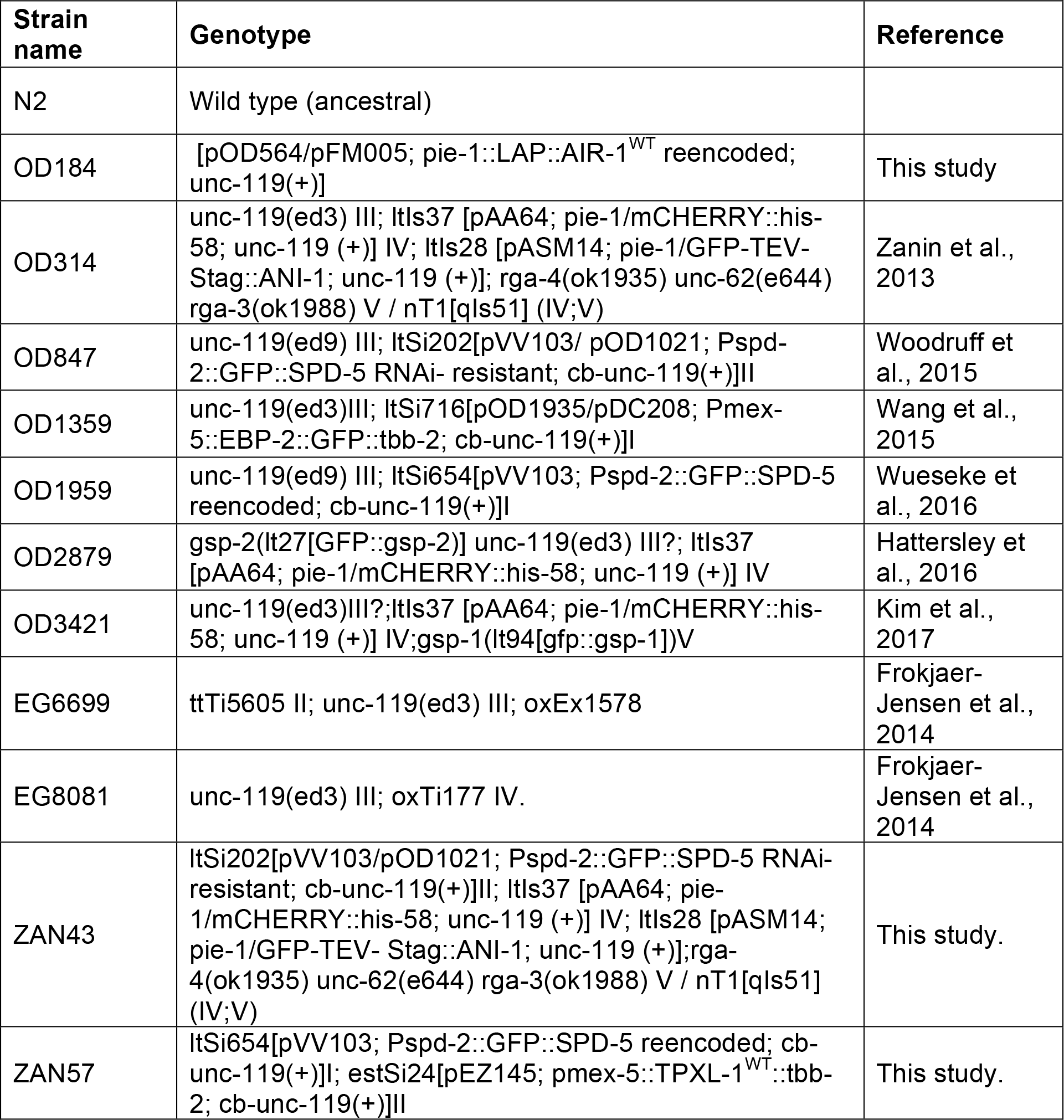
*C. elegans* strains

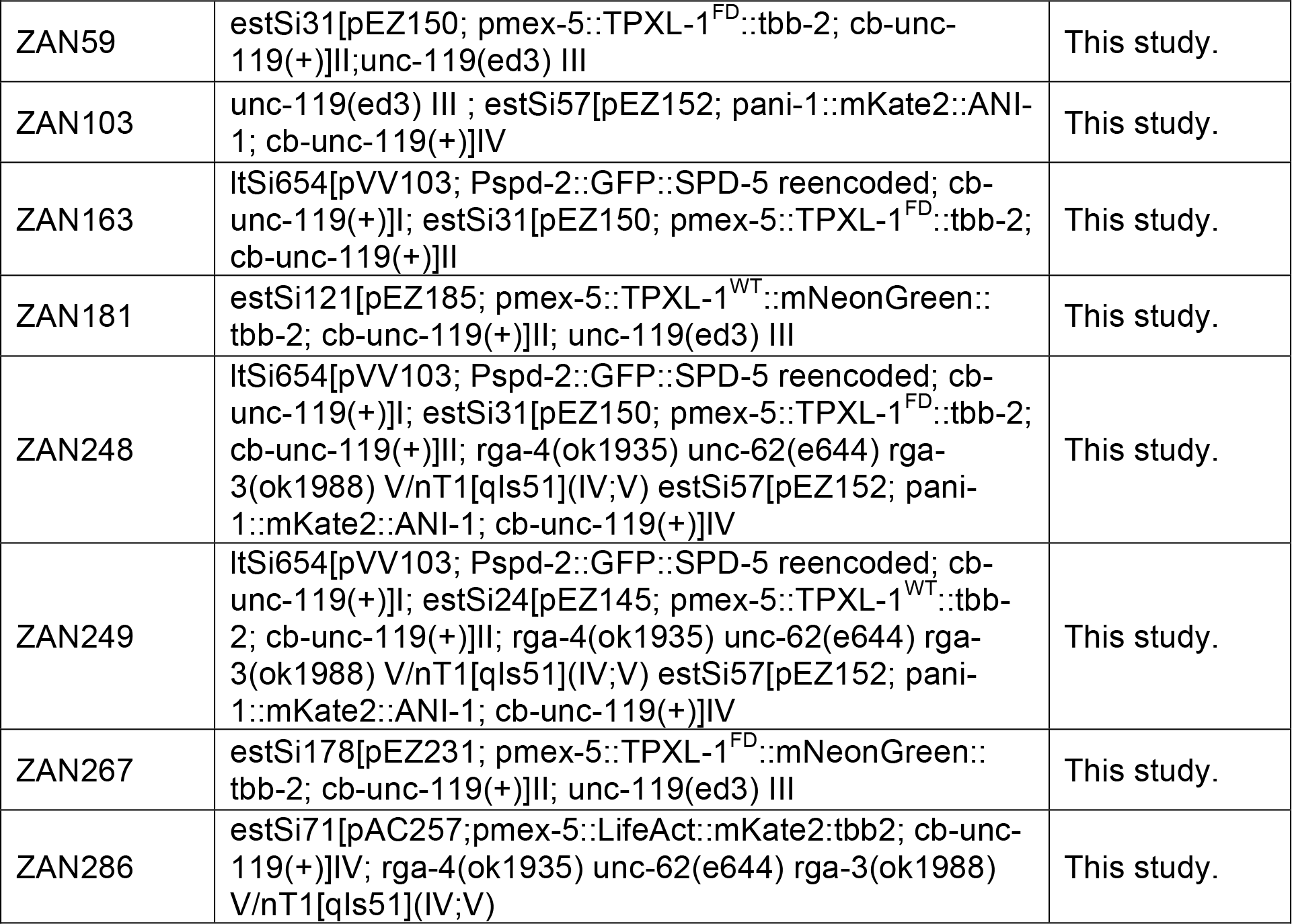

**Table 2.**
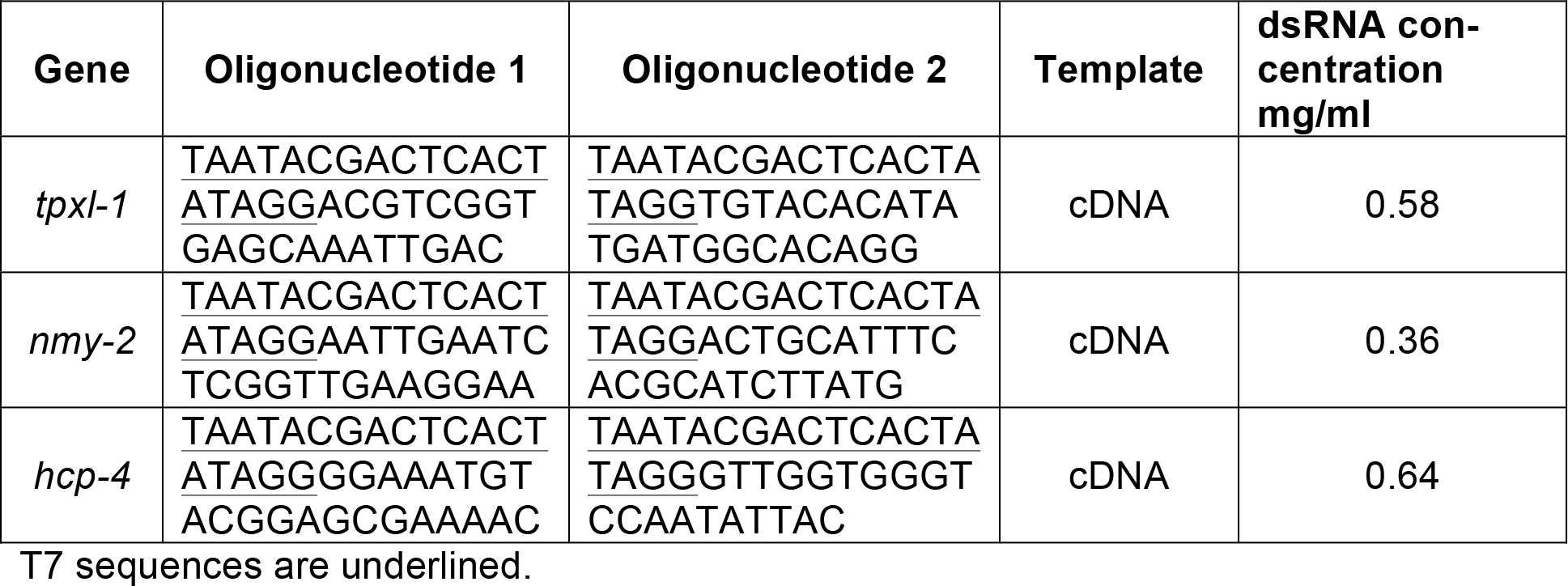
dsRNA production for *C. elegans*

## ACKNOWLEDGMENTS

We thank A. Desai and B. Conradt for critical comments on the manuscript. This work was supported by an NIH grant to K.O. (GM074207), the DFG Emmy-Noether-Program to E.Z. (ZA619/3-1); T.K. was supported by an NIH grant to A. Desai (GM074215). Some strains were provided by the CGC, which is funded by NIH Office of Research Infrastructure Programs (P40 OD010440). We thank H. Bringmann for the mKate2 plasmid, A.A. Hyman for the TPXL-1 antibody and the Leonhardt and Conradt lab for sharing reagents and expertise. We thank T. Mikeladze-Dvali for fruitful discussions, H. Harz for outstanding support with the microscopy and L. Jocham, N. Lebedeva and M. Schwarz for excellent technical support. K.O. receives salary and other support from the Ludwig Institute for Cancer Research. S.M. is a member of IMPRS-LS (International Max Planck Research School for Molecular Life Sciences) and J.S. is a member of the LSM (Life Science Munich) graduate program and both thank their programs for support.

## Online supplemental material

**Figure S1**, which is related to Figure 4 and 5, shows the transgenes encoding wild-type untagged and mNG-tagged TPXL-1 and their ability to rescue embryonic lethality resulting from endogenous TPXL-1 depletion.

**Figure S2**, related to Figure 4, shows the localization of endogenous TPXL-1 and Aurora A^AIR-1^ to astral microtubules in anaphase and measurement of microtubule growth and nucleation rate in control embryos at metaphase and early and late anaphase.

**Movie S1, related to Figure 1**

**Movie S2, related to Figure 1**

**Movie S3, related to Figure 2**

**Movie S4, related to Figure 3**

**Movie S5, related to Figure 4**

**Movie S6, related to Figure 4**

**Movie S7, related to Figure 5**

## SUPPLEMENTARY FIGURE LEGENDS

**Figure S1, related to Figure 4 and 5:**
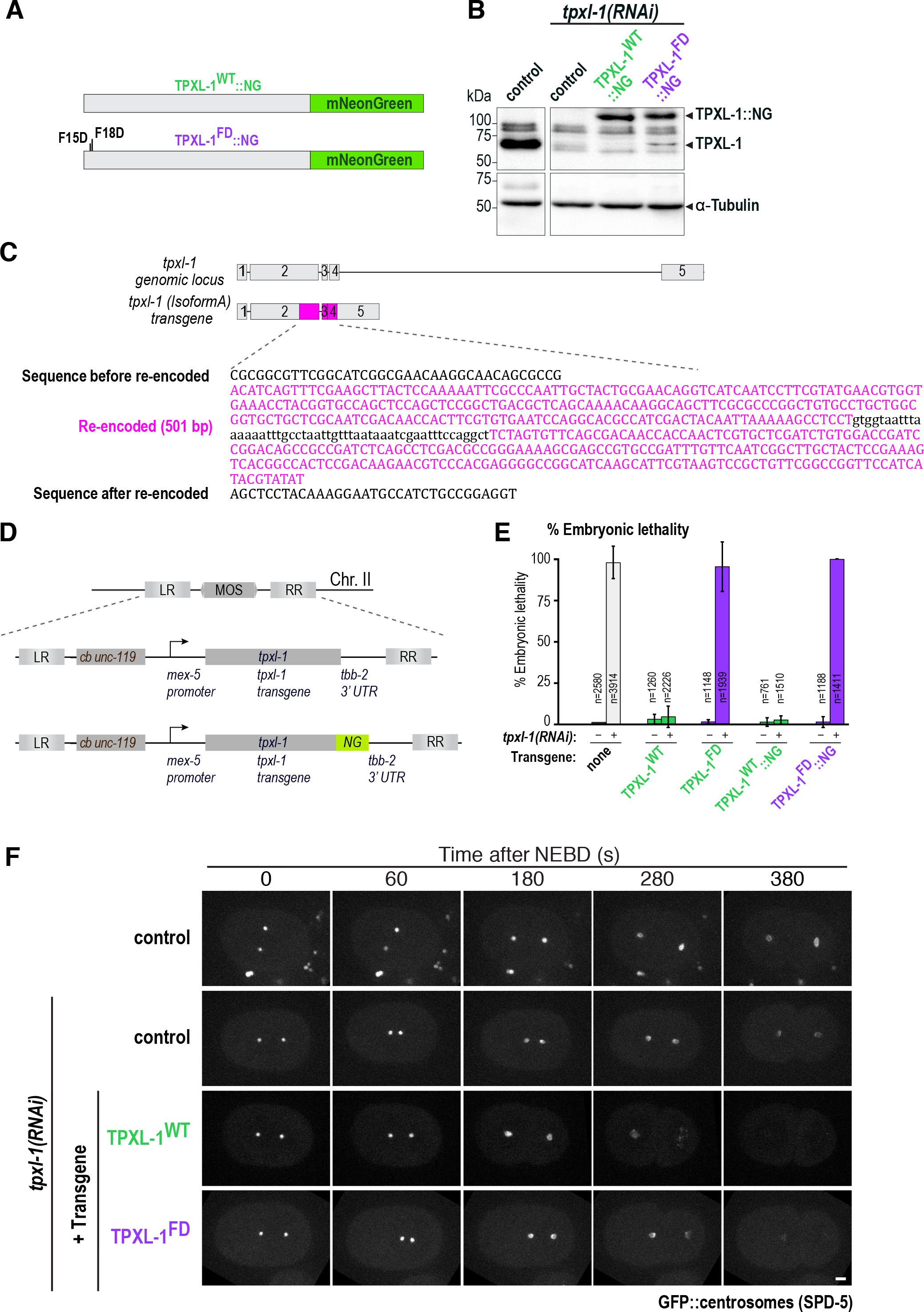
Transgenes encoding wild-type untagged and mNeonGreen tagged TPXL-1 rescue embryonic viability resulting from endogenous TPXL-1 depletion. (**A**) Schematics showing TPXL-1^WT^ or TPXL-1^FD^ tagged with NeonGreen. (**B**) Immunoblots of control (N2) and transgenic animals expressing RNAi-resistant TPXL-1^WT^::NG or TPXL-1^FD^::NG. Endogenous TPXL-1 was depleted by RNAi as indicated and immunoblots were probed for TPXL-1 and α-tubulin as a loading control. (**C**) Schematic representation of intron-exon organization of *C. elegans* TPXL-1. To make the TPXL-1 transgene RNAi resistant, a region including exon 3 and parts of exons 2 and 4 was re-encoded. The intron between exon 2 and 3 was maintained. (**D**) The untagged and NG-tagged TPXL-1 transgenes, with the *mex-5* promoter and *tbb-2* 3’UTR, were integrated into chromosome II using MosSCI (Frøkjær-Jensen et al., 2008). (**E**) Graph plotting percent embryonic lethality for the indicated conditions. Error bars are SD and n = number of embryos analyzed. (**F**) Maximum intensity projections of 5 confocal planes (1.5 μm apart) of embryos expressing GFP::SPD-5 either with TPXL-1^WT^, TPXL-1^FD^ or without a transgene (control). Endogenous TPXL-1 was depleted by RNAi. The distance between the two centrosomea was measured over time and is quantified in Fig. 5D. Time after NEBD is indicated and scale bar is 5μm.

**Figure S2, related to Figure 4:**
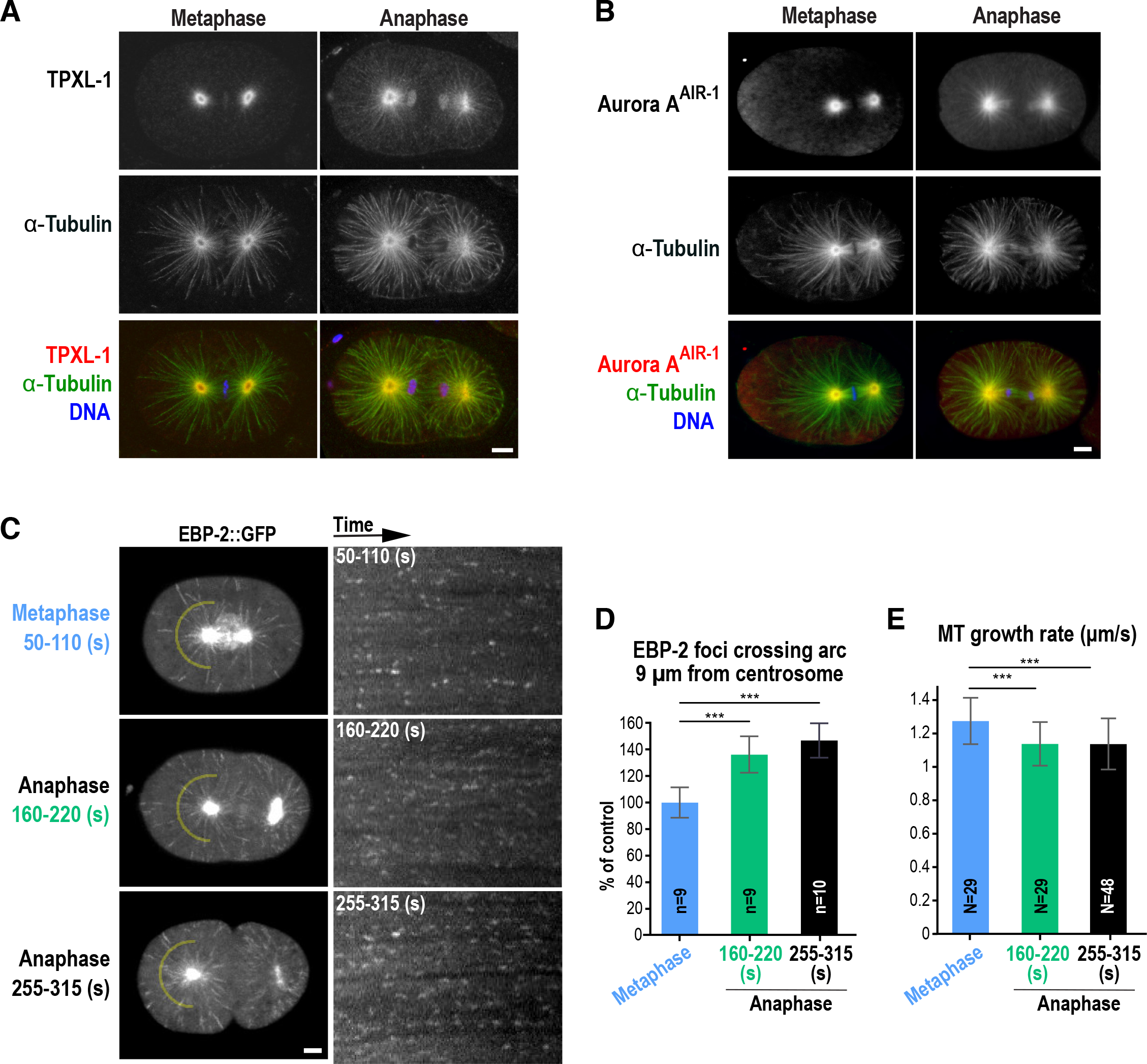
Endogenous TPXL-1 and Aurora A^AIR-1^ localizes to astral microtubules in anaphase. (**A**) Confocal images of fixed metaphase and anaphase embryos stained for endogenous TPXL-1, α-tubulin, and DNA (prometaphase & metaphase n=8, anaphase n=9 embryos). To visualize astral microtubules without saturating the aster centers, a gamma of 2 was used for all images, which were scaled identically. **(B)** Confocal images of fixed metaphase (n=5) and anaphase (n=5) embryos stained for Aurora A^AIR-1^, α-tubulin, and DNA. To visualize astral microtubules without saturating the aster centers, a gamma of 1.5 was used for all images, which were scaled identically. (**C**) Representative projections of the images acquired every 400 ms over a 4s interval in metaphase (n=9), early anaphase (160-220s after NEBD; n=9) and late anaphase (255-315s after NEBD; n=10) control embryos expressing EBP-2::GFP. Kymographs generated for the indicated conditions (***right***) were used to count the number of EBP-2::GFP foci that crossed an arc 9 μm away from the anterior centrosome (***yellow***). (**D,E**) Graphs plot the number of EBP-2 foci crossing an arc 9 μm from the centrosome (D) and the microtubule growth rates (E) at the indicated times in control embryos. (n=number of embryos and error bars are SD in (D); N=number of microtubules tracked in 3 or more embryos per condition and error bars are SD in (E)). All scale bars are 5μm.

## MOVIE LEGENDS

**Movie S1, related to Figure 1:** *C. elegans* one-cell embryos expressing GFP::anillin (***cyan***) and mCherry::histone (***red***) without (control, ***left***) and with (***right***) myosin depletion. Images were acquired every 20s on an UltraVIEW VoX spinning disk confocal microscope (PerkinElmer) attached to an Axio Observer D1 stand (Carl Zeiss, Inc.), equipped with a 63× 1.4 NA Plan-Apochromat oil immersion objective and a Hamamatsu EMCCD C9100-50 camera (1000x1000 pixel). Movie starts 60s after NEBD. Playback rate is 60× real time (3 frames/second).

**Movie S2, related to Figure 1:** Two representative myosin depleted & *rga-3/4*Δ embryos expressing GFP::anillin (***cyan***) and mCherry::histone (***red***). Images were acquired every 20s on an UltraVIEW VoX spinning disk confocal microscope (PerkinElmer) attached to an Axio Observer D1 stand (Carl Zeiss, Inc.), equipped with a 63x 1.4 NA Plan-Apochromat oil immersion objective and a Hamamatsu EMCCD C9100-50 camera (1000x1000 pixel). Movie starts 60s after NEBD. Playback rate is 60× real time (3 frames/second).

**Movie S3, related to Figure 2:** Myosin depleted & *rga-3/4Δ C. elegans* embryos without (***left***) and with *hcp-4(RNAi)* (***right***) expressing GFP::anillin and a GFP-tagged centrosome marker (***cyan***) along with mCherry::histone (***red***). Images were acquired every 20s on an UltraVIEW VoX spinning disk confocal microscope (PerkinElmer) attached to an Axio Observer D1 stand (Carl Zeiss, Inc.), equipped with a 63× 1.4 NA Plan-Apochromat oil immersion objective and a Hamamatsu EMCCD C9100-50 camera (1000x1000 pixel). Playback rate is 60× real time (3 frames/second). Movie starts 180s after NEBD.

**Movie S4, related to Figure 3:** Myosin depleted & *rga-3/4Δ C. elegans* embryos expressing GFP::anillin and a GFP-tagged centrosome marker (***cyan***) along with mCherry::histone (***red***). TPXL-1 (***left***) or TPXL-1 & HCP-4 (***right***) were additionally depleted by RNAi. Images were acquired every 20s on an UltraVIEW VoX spinning disk confocal microscope (PerkinElmer) attached to an Axio Observer D1 stand (Carl Zeiss, Inc.), equipped with a 63× 1.4 NA Plan-Apochromat oil immersion objective and a Hamamatsu EMCCD C9100-50 camera (1000x1000 pixel). Playback rate is 60x real time (3 frames/second). Movie starts 180s after NEBD.

**Movie S5, related to Figure 4:** Representative examples of *C. elegans* embryos expressing TPXL-1^WT^::NG (***left***) or TPXL-1^FD^::NG (***right***) from RNAi-resistant transgenes after depletion of endogenous TPXL-1 by RNAi. Images were acquired on a Nikon eclipse Ti spinning disk confocal controlled by NIS Elements 4.51 software and equipped with a 100x 1.45 NA Plan-Apochromat oil immersion objective and Andor DU-888 X11056 camera. Elapsed time and anaphase onset are indicated.

**Movie S6, related to Figure 4:** Representative examples of *C. elegans* embryos expressing GFP::Aurora A^AIR-1^ without(***left***) and with *tpxl-1(RNAi)* (***right***). Images were acquired on a Nikon eclipe Ti spinning disk confocal controlled by NIS Elements 4.51 software and equipped with a 100x 1.45 NA Plan-Apochromat oil immersion objective and Andor DU-888 X11056 camera. Elapsed time and anaphase onset are indicated.

**Movie S7, related to Figure 5:** Representative myosin depleted & *rga-3/4Δ C. elegans* embryos expressing TPXL-1^WT^ (***left***) or TPXL-1^FD^ (***right***) together with mKate2::anillin (***magenta***). Embryos were additionally depleted of endogenous TPXL-1 and HCP-4 by RNAi. Images were acquired every 20s on an UltraVIEW VoX spinning disk confocal microscope (PerkinElmer) attached to an Axio Observer D1 stand (Carl Zeiss, Inc.), equipped with a 63× 1.4 NA Plan-Apochromat oil immersion objective and a Hamamatsu EMCCD C9100-50 camera (1000×1000 pixel). Playback rate is 60x real time (3 frames/second). Movie starts 180s after NEBD.

